# High Resolution Class I *HLA -A, -B*, and -*C* Diversity in Eastern and Southern African Populations

**DOI:** 10.1101/2024.09.04.611164

**Authors:** Alabi W. Banjoko, Tiza Ng’uni, Nitalia Naidoo, Veron Ramsuran, Olivier Hyrien, Zaza M. Ndhlovu

## Abstract

Africa remains significantly underrepresented in high-resolution Human Leukocyte Antigen (HLA) data, despite being one of the most genetically diverse regions in the world. This critical gap in genetic information poses a substantial barrier to HLA-based research on the continent. In this study, Class I HLA data from Eastern and Southern African populations were analysed to assess genetic diversity across the region. We examined allele and haplotype frequency distributions, deviations from Hardy-Weinberg Equilibrium (HWE), linkage disequilibrium (LD), and conducted neutrality tests of homozygosity across various populations. Additionally, the African HLA data were compared to those of Caucasian and African American populations using the Jaccard index and multidimensional scaling (MDS) methods. The study revealed that South African populations exhibited 50.4% more genetic diversity within the Class I HLA region compared to other African populations. Zambia showed an estimated 36.5% genetic diversity, with Kenya, Rwanda and Uganda showing 35.7%, 34.2%, and 31.1%, respectively. Furthermore, an analysis of in-country diversity among different tribes indicated an average Class I HLA diversity of 25.7% in Kenya, 17% in Rwanda, 2.8% in South Africa, 13.6% in Uganda, and 6.5% in Zambia. The study also highlighted the genetic distinctness of Caucasian and African American populations compared to African populations. Notably, the differential frequencies of disease- promoting and disease-preventing HLA alleles across these populations emphasize the urgent need to generate high-quality HLA data for all regions of Africa and its major ethnic groups. Such efforts will be crucial in enhancing healthcare outcomes across the continent.

**Author Summary:** This study investigated the diversity of class I HLA in the eastern and southern regions of the African continent using a population genetics approach. Analysis of HLA data at both country and tribal levels revealed significant genetic differences and the unique characteristics of these populations compared to Caucasian and African American populations in the United States. The differential frequencies of disease-promoting and disease-preventing HLA alleles across these populations suggest that large-scale vaccine administration may be ineffective without a thorough understanding of the HLA composition of each population. This study highlights the urgent need to generate high-quality HLA data across all regions of Africa and its major ethnic groups. Such comprehensive data collection is essential for optimizing vaccine design, deepening our understanding of HLA-disease associations, and ultimately improving healthcare outcomes across the continent.

## INTRODUCTION

The Human Leukocyte Antigen (HLA) complex consists of highly polymorphic genes that code for surface proteins responsible for presenting antigens to T cells as part of an immune response to infections [1]. According to the IPD-IMGT/HLA database, more than 40,000 HLA alleles have been identified and the total HLA allele variation is estimated to be several millions across the different populations around the world [2]. Africa, often referred to as the cradle of humankind [3, 4], boasts of the highest levels of human genetic diversity in the world [5]. This rich genetic diversity is a result of the continent’s long evolutionary history, complex demographic processes and genetic admixture that have shaped its populations over time [4, 6]. However, population data sets on some of the databases such as Allele Frequency Net Database (AFND) which provides the scientific community with a freely available repository for the storage of frequency data including alleles, genes, haplotypes, and genotypes have reported very limited HLA frequency data for African populations [7–9]. Moreover, many ethnic populations in Sub-Saharan Africa are underrepresented in medical genomics studies due to limited research, particularly on *HLA* alleles, compared to developed countries [10]. This discrepancy in *HLA* allele data is also reflected in the IPD-IMGT/HLA database, where most submissions originate from Europe, America, and Australia (IMGT/HLA Database, released of July 2024, <u>IPD-IMGT/HLA Database</u> <u>(ebi.ac.uk)</u>). This indicates a significant lack of HLA typing infrastructure in Sub-Saharan Africa, further contributing to the scarcity of HLA data for these populations.

The *HLA* genes have been widely studied over the years due to their extensive allelic variability across diverse populations and their importance in host immune responses, therapy and organ transplantation [2, 8, 11–13]. In addition, some *HLA* alleles have been associated with either protection against or susceptibility to a wide range of autoimmune and infectious diseases as well as drug-induced hypersensitivity and cancer [14]. For instance, HLA class I alleles *such HLA- B*27, HLA-B*52, HLA-B*57* and *HLA-B*81* have been linked to protection against HIV disease progression (protective alleles) whereas *HLA-B*35, HLA-B*51:01*, and *HLA-B*58:02* have been linked to rapid disease progression (disease-susceptible alleles) [15, 16]. However, HLA alleles and haplotypes do not occur at the same frequency in different populations. For example, in Caucasians, *HLA-B*58:02* which is linked to HIV disease susceptibility is mainly absent whereas it is highly prevalent in the African population [15]. Similarly, the protective allele *HLA-B*57:01* is highly prevalent in the Caucasian population whereas it is largely absent in the African population [15, 17, 18].

Leveraging HLA diversity data can lead to more tailored therapies and inform the rational design of T cell-based vaccines that will be efficacious across different populations. In this study, population genetics approaches have been used for understanding HLA (genetic) diversity in the eastern and southern African regions. The study provides an insight into the extensive diversity of the allelic and haplotype frequencies within five African populations and compared to the Caucasian and African American populations.

## RESULTS

### Genetic Diversity Between African and U.S. Populations

To address the extent of HLA differences between the African populations and the US populations, we compared HLA data from the African sub-regions to the Caucasian and the African American populations. Although, the Caucasian and African American HLA studies have received some attention in literature [19–21], this study demonstrated that HLA data from the US populations cannot be a true representative of the African HLA population data. We computed allele frequencies across all populations in this study to help identify complex genetic traits and discover HLA disease associations [22, 23]. Frequencies of alleles were estimated by direct counting. The full list of alleles and their frequencies across populations is detailed in **Supplementary Tables 1, 2 and 3**. Allelic frequency distributions vary across populations. Some alleles frequencies are either high or low, while others may be present or absent across populations. Allele frequencies (for countries) were sorted in descending order within each population and alleles with frequencies of at least 5% were plotted and presented in Figure 1 for all loci.

**Figure 1.**
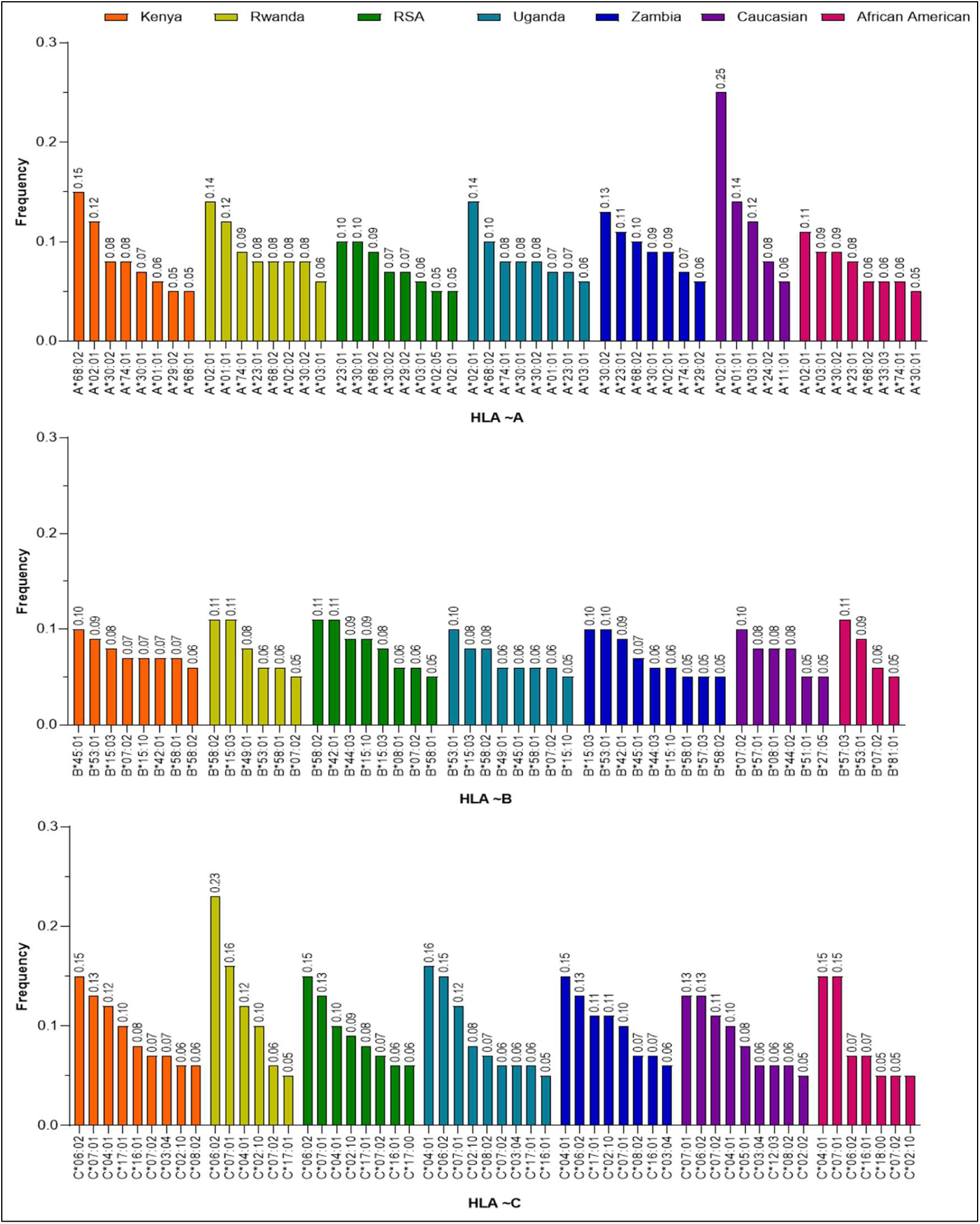
Most frequent (≥ 5%) HLA alleles within each population. Distinction in allele frequencies testify to HLA genetic architectural diversity among the populations. HLA-B has relatively less frequencies.

### HLA-A

*HLA-A*02:01* is present among the top 5% alleles in all the populations at different high frequency level with it being highest in Caucasians at 25% frequency. However, this allele was relatively low in the South African (RSA) population with a frequency of 5.1% as presented in Figure 1. Also, *HLA-A*68:02* was observed at a relatively high frequency (> 8%) in the African populations but low (6%) in AFAM. Interestingly, this allele was not among the top 5% in the Caucasian population. *HLA-A*68:01, A*02:02, A*02:05*, and *A*11:01* were observed among the top 5% frequent alleles and only found in Kenya (5%), Rwanda (8%), RSA (5%), and Caucasian (6.4%) populations respectively (Figure 1 and Supplementary Table 1).

### HLA-B

It was observed that *HLA-B*15:03, HLA-B*58:01* and *HLA-B*58:02* were present among the top 5% alleles in all populations (though at different frequency levels) except the US populations (Figure 1). Also, *HLA-B*57:03* was only present in AFAM (10.8%) and Zambia (5.3%) among the top 5% frequent alleles. *HLA-B*27:05* and *HLA-B*57:01* were only present in the Caucasian population whereas, *HLA-B*81:01* was only present in the AFAM population among the top 5% frequent alleles. *HLA B*07:02* was among the top 5% frequent alleles in all populations except in Zambia (Figure 1, Supplementary Table 2).

### HLA-C

In locus C, *HLA-C*04:01*, *HLA-C*06:02* and *HLA-C*07:01* were highly observed and listed among the top five of the 5% most frequent alleles in all the populations (Figure 1). However, *HLA-C*02:10* was among the top 5% frequent alleles in the African and AFAM populations but was not among the top 5% alleles in the Caucasian population. Also, *HLA-C*17:01* was among the top 5% alleles only in the African populations. In the Caucasian population, *HLA-C*05:01* and *HLA-C*12:03* were present at 8% and 6% respectively but were not among the top 5% frequent alleles in other populations (Figure 1, Supplementary Table 3).

Jaccard index was used to quantity the similarity (or dissimilarity) between two populations in terms of alleles composition and genetic makeup. The Jaccard index was obtained by determining the alleles that are simultaneously present in two populations. The structure of the alleles was then used to determine the Jaccard similarity indices, converted into percentages and drawn as a non-clustered heat map. The darker the red colour in the heatmap, then the more similar two corresponding populations. Generally, Figure 2C shows low level of genetic similarities between the African and US populations. The Caucasian population had the lowest similarity index to the African populations in all the HLA locus considered. Similarly, the African American population showed relatively higher similarity indices to the African populations than the Caucasian population in all loci.

**Figure 2.**
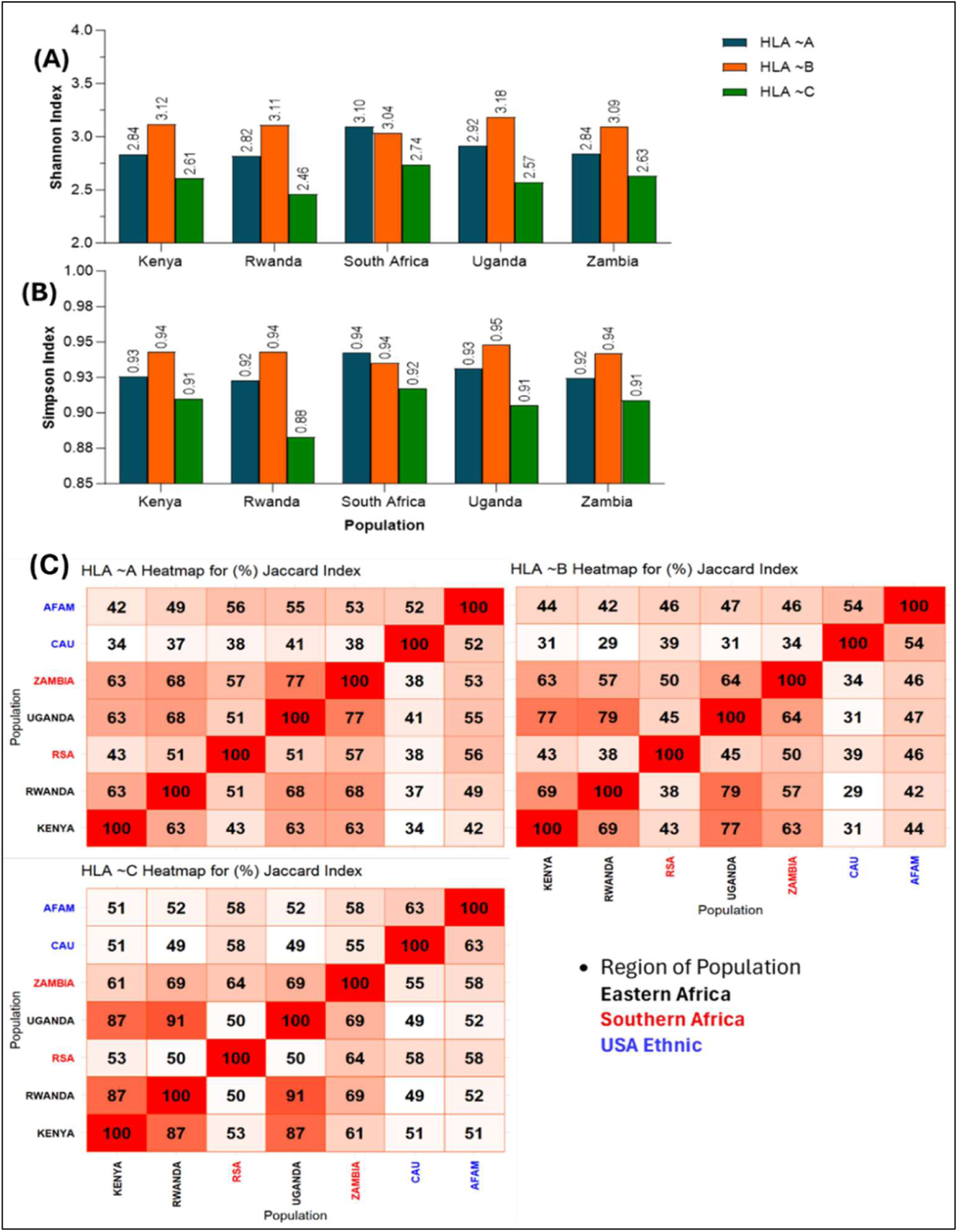
Graphs of Shannon **(A)**, Simpson **(B)** indices across African populations and **(C)** non-clustered heatmap of similarity index (Jaccard) among populations. A and B explain the in-country diversity. The higher the index values the more the diversity of the population at that locus. C quantify the genetic similarities (in %) among populations. The darker the red colour, the more similar the two populations involved.

In addition to the individual *HLA* alleles, the study determined the extent to which haplotypes (specific combination of alleles inherited together on the same chromosome) overlap between populations. This was determined using MDS to visualize the genetic distances (cartograph) at all the haplotype loci. Analysis was carried out on the relative frequency of haplotypes in each population relative to other populations. Haplotype frequencies from each population data were dimensionally reduced using MDS to create a 2-dimensional genetic cartograph. Based on the analysis, two countries are close to each other on the map if the distribution of the haplotypes in these two populations are close to each other, relative to the distribution observed in the other countries. In Figure 3, the African American population is relatively closer to the African populations compared to the Caucasian which is farther away from the African populations at the global haplotypes.

**Figure 3.**
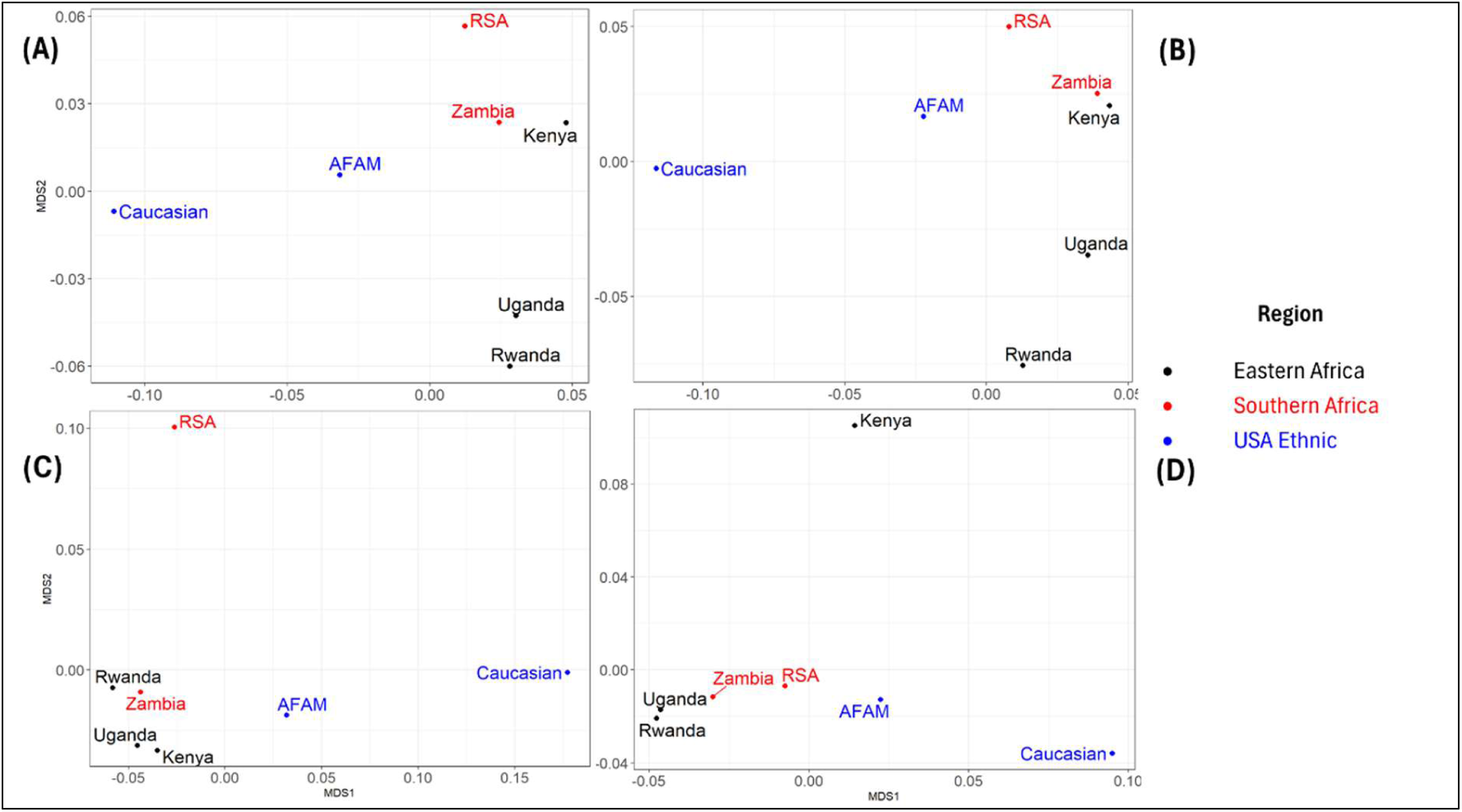
Cartography of the genetic distance in global haplotypes between populations. **(A), (B), (C)** and **(D)** represents haplotype A∼B, A∼C, B∼C, and A∼B∼C respectively. The figure visualizes the genetic distance between African and US ethic populations.

### Genetic Diversity Within African Populations

To better define HLA genetic diversities among African populations at both country and tribal levels, we computed within populations diversities using the Shannon and Simpson diversity indices. Similarly, the Jaccard index in Figure 2C present a diversity index to compare the HLA differences between African populations. The Shannon and Simpson indices were determined at each locus and across African populations (**see Supplementary Figure 1 for tribes**). Figure 2A provides a summary of the Shannon index which accounts for alleles specie richness and evenness of their abundance. Also, Figure 2B provides a summary of Simpson indices which account for probability that two alleles taken from the sample at random are of different types. In Figures 2A and 2B, all populations present the natural polymorphic structures of the HLA alleles except for South Africa wherein HLA-A is slightly more evenly distributed than HLA-B. Generally, the higher values observed in all populations at different loci indicate high level of genetic diversity within each of the African populations. A similar trend of results were observed in the tribal populations of each country in the African sub region (**Supplementary Figure 1A and 1B**).

In Figure 2C, the highest values of the Jaccard index among the eastern African countries were observed between Rwanda and Uganda with similarity indices of 68%, 79% and 91% at locus A, B and C respectively. South Africa and Zambia had similarity indices of 57%, 50% and 64% at locus A, B and C respectively. Interestingly, Uganda and Zambia had the highest similarity index of 77% at locus A. Conversely, South Africa had the lowest similarities with other African countries at all loci except with Zambia at locus C where the similarity index was relatively higher (64%). At the tribal level (**Supplementary Figure 1C**), the Jaccard index also shows various values of similarity indices across tribes within the African countries. The Zulu tribe show low (24% ≤ *JI* ≤ 56%) similarity to other African tribes at all loci except with Bemba tribe (62% at locus C) from Zambia. It was observed that only few tribes from some of the African countries have similarity above 80%. For instance, a high similarity of 82% was observed between the Nsenga tribe of Zambia and Munyankole tribe in Uganda at locus A.

Also the cartograph shows that at loci B:C, South Africa (RSA) was observed to be farther away from other African countries (Figure 3C). Similarly, Kenya is seen to be far from the rest of the African countries at loci A:B:C (Figure 3D). Zambia and Kenya were observed to be closer at loci A:B and A:C (Figures 3A and 3B), while Uganda and Rwanda were closer at loci A:B and A:B:C (Figures 3A and 3D). We also observed close genetic distances between Zambia and Rwanda as well as Uganda and Kenya at loci B:C (Figure 3C), whereas, Uganda and Rwanda were closer at loci A:B:C (Figure 3D). Furthermore, the cartograph at tribal level as presented in **Supplementary Figure 2** shows that the genetic distances between the Kenyan tribes (Kikuyu and Luhya) are farther apart from each other at all loci. The Ugandan tribes in this study (Muganda, Munyankole and Munyarwanda) were observed to be far apart from each other at all loci except for loci B:C where they seem to be relatively closer. Also, the Lozi tribe in Zambia did not exhibit the same genetic closeness as other Zambian tribes at all loci (**Supplementary Figures 2**). The Zulu tribe from South Africa exhibited genetic closeness to tribes in Zambia and Uganda at all loci except B:C where it was relatively farther from other African tribes.

The rarefaction curves in Figure 4 were employed to evaluate completeness of samples and uniqueness of alleles for each population (country and tribe) as a function of the number of participants. This was determined by creating a subsample of size *n* and counting the number of unique HLA alleles included in the subsample for any given (HLA alleles, population) pair. Subsampling was done at random without replacement and was repeated for different values of *n* = 1, . . ., *N*_p_, where *N*_p_ denotes the number of participants from the selected population. We next plotted the number of unique HLA alleles as a function of the number of participants by population. The overall numbers and percentage frequencies of alleles observed across populations at each locus are presented in **Table 1**.

**Figure 4.**
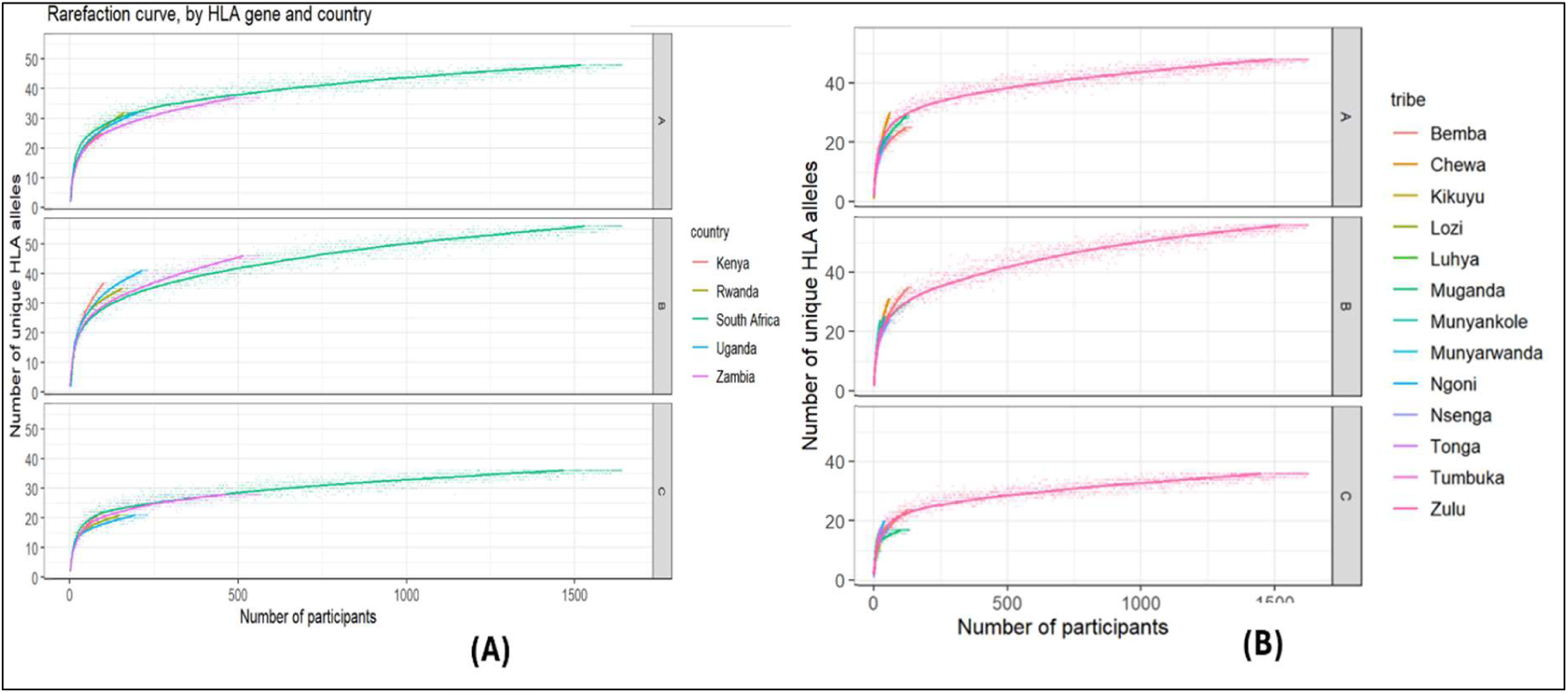
Rarefaction curves by HLA gene and populations estimating the allelic diversity or richness. It explains detection of more allelic variants at each locus as more participants are selected in each population.

**Table 1.** Number of alleles for Class I alleles according to country of subjects.

| Populations | No of participants | No of Alleles |  |  | Frequency of Alleles (%) |  |  |
| --- | --- | --- | --- | --- | --- | --- | --- |
|  |  | HLA ~A | HLA ~B | HLA ~C | HLA ~A | HLA ~B | HLA ~C |
| Kenya | 109 | 25 | 37 | 22 | 22.9 | 33.9 | 20.2 |
| Rwanda | 173 | 32 | 35 | 21 | 18.5 | 20.2 | 12.1 |
| South Africa | 1640 | 48 | 56 | 36 | 2.9 | 3.4 | 2.2 |
| Uganda | 231 | 32 | 41 | 21 | 13.9 | 17.7 | 9.1 |
| Zambia | 565 | 37 | 46 | 28 | 6.5 | 8.1 | 5.0 |
| Caucasian | 1765 | 61 | 101 | 40 | 3.5 | 5.7 | 2.3 |
| African American | 661 | 50 | 78 | 40 | 7.6 | 11.8 | 6.1 |

The rarefaction curves in Figure 4 indicate that the curves typically rises quickly initially as unique alleles were observed then levelled of as only few rare alleles remain to be observed. Also, the curves presented the natural diversity structures of alleles in each locus [24]. HLA-B had the highest allelic diversity in all populations (countries and tribes), followed by HLA-A, while HLA-C had the least allelic diversity. Visual inspection of the curves also supports the genetic diversities observed in Table 1 across the African countries. In Table 1, Kenya is observed to have the highest allelic diversity at all loci, followed by Rwanda, then Uganda and Zambia, while South Africa has the least diversity at all loci among the African countries.

### HLA Alleles Linked to Immune Responses and Disease Outcomes

Additionally, we investigated immune/disease associations of some known alleles at each locus to determine the important differences in the HLA alleles that are associated with immune/disease outcomes for the different populations. This will assist in understanding the genetic basis of immune responses and the association with diseases such as HIV, leading to improved diagnostics, treatments, and preventive strategies [25, 26]. According to literature, some alleles are grouped as either Bw4 or Bw6 [27]. The Bw4 and Bw6 are epitopes found on most HLA-B and few HLA-A proteins, which play a role in immune responses [28]. In addition, other alleles are classified as either protective or disease susceptible based on different populations and studies [27–31]. This study observed the frequencies of the known immune and HIV disease associated alleles in each of the populations and the results are presented in Table 2. It was observed that at locus A and B, Bw4 group alleles were either observed at an extremely low frequency or not observed at all in all the populations. *HLA-A*74:01* (protective allele) was observe at relatively higher frequencies in Kenya, Rwanda, Uganda, Zambia and African American than in South Africa and very low in the Caucasian. Also, *HLA-A*25:01* was not observed in all the African populations and had a very low frequency in the African American population. The HIV disease-susceptible allele at locus A (*A*36:01*) was observed at a relatively low frequency in South African and Caucasian populations compared to other populations. Similarly, the Bw4 and Bw6 alleles were technically not observed in the African populations at locus B. Although *HLA-B*39:01* was observed in the Zambia population but at an extremely low frequency. Among the protective alleles, *HLA-B*27:05* was observed at a low frequency in South Africa among the African populations compared to the Caucasian and African American populations. Conversely, *HLA-B*42:01* and *B*44:03* were observed at the high frequencies in both South African and Zambian populations. *HLA-B*52:01* was only observed in Kenya and South Africa among the African populations with low frequencies. While *HLA-B*57:01* was observed at extremely frequencies in African populations, it was observed at a high frequency (7.9%) in the Caucasian population. Similarly, among the HIV disease-susceptible alleles at locus B, frequencies of *HLA-B*35:01*, *B*35:02* and *B*35:03* were generally low in African populations and specifically lower in the South African population. In African populations, *HLA- B*07:02* was observed at a relatively high frequency in Kenya compared to other African countries while the Caucasian population had the highest frequency of *B*07:02* among all populations. Also, among the HIV disease-susceptible alleles at locus B, *HLA-B*08:01* was observed at a relatively high frequency in the Caucasian population compared to other populations. Similarly, this allele was at a high frequency in South Africa compared to other African populations. Interestingly, *HLA-B*58:02* was observed at higher frequencies in the African populations compared to the US populations. This allele seems to have very high frequency in South Africa (11.4%) and Rwanda (11.3%) compared to other African countries.

**Table 2.** Grouping of alleles and allele frequencies among populations at different locus.

| Locus | Groups | HLA | Population |  |  |  |  |  |  |
| --- | --- | --- | --- | --- | --- | --- | --- | --- | --- |
|  |  |  | Kenya | Rwanda | South Africa | Uganda | Zambia | Caucasian | African American |
| A | BW4 | A*24:03 |  |  | 0.0003 |  |  | 0.0037 | 0.0023 |
|  | Protective | A*25:01 |  |  |  |  |  | 0.0292 | 0.0061 |
|  |  | A*32:01 |  | 0.0029 | 0.0043 | 0.0152 | 0.0035 | 0.0462 | 0.0166 |
|  |  | A*74:01 | 0.0780 | 0.0896 | 0.0366 | 0.0844 | 0.0690 | 0.0006 | 0.0575 |
|  | HIV Disease-Susceptible | A*36:01 | 0.0275 | 0.0318 | 0.0043 | 0.0433 | 0.0460 | 0.0014 | 0.0212 |
| B | BW4 | B*51:02 |  |  |  |  |  |  | 0.0008 |
|  | BW6 | B*39:01 |  |  |  |  | 0.0009 | 0.0105 | 0.0045 |
|  |  | B*39:02 |  |  |  |  |  | 0.0003 |  |
|  | Protective | B*13:02 | 0.0138 | 0.0145 | 0.0162 | 0.0108 | 0.0106 | 0.0283 | 0.0129 |
|  |  | B*14:02 | 0.0321 | 0.0318 | 0.0125 | 0.0346 | 0.0327 | 0.0456 | 0.0272 |
|  |  | B*27:05 |  |  | 0.0015 |  |  | 0.0501 | 0.0144 |
|  |  | B*42:01 | 0.0734 | 0.0405 | 0.1079 | 0.0433 | 0.0912 | 0.0011 | 0.0378 |
|  |  | B*44:03 | 0.0275 | 0.0347 | 0.0918 | 0.0216 | 0.0646 | 0.0473 | 0.0484 |
|  |  | B*52:01 | 0.0046 |  | 0.0003 |  |  | 0.0193 | 0.0197 |
|  |  | B*57:01 |  |  | 0.0003 | 0.0022 | 0.0018 | 0.0793 | 0.0106 |
|  |  | B*57:02 | 0.0092 |  | 0.0079 | 0.0130 | 0.0097 | 0.0014 | 0.0129 |
|  |  | B*57:03 | 0.0413 | 0.0347 | 0.0229 | 0.0390 | 0.0531 | 0.0105 | 0.1082 |
|  |  | B*58:01 | 0.0688 | 0.0607 | 0.0503 | 0.0584 | 0.0549 | 0.0133 | 0.0416 |
|  |  | B*81:01 | 0.0183 | 0.0260 | 0.0418 | 0.0433 | 0.0257 | 0.0006 | 0.0522 |
|  | HIV Disease-Susceptible | B*07:02 | 0.0734 | 0.0549 | 0.0561 | 0.0563 | 0.0336 | 0.0952 | 0.0552 |
|  |  | B*08:01 | 0.0046 | 0.0173 | 0.0616 | 0.0238 | 0.0301 | 0.0756 | 0.0371 |
|  |  | B*18:01 | 0.0183 | 0.0289 | 0.0354 | 0.0346 | 0.0319 | 0.0428 | 0.0242 |
|  |  | B*35:01 | 0.0321 | 0.0318 | 0.0177 | 0.0238 | 0.0354 | 0.0456 | 0.0386 |
|  |  | B*35:02 |  | 0.0029 | 0.0015 | 0.0022 | 0.0009 | 0.0079 | 0.0015 |
|  |  | B*35:03 |  |  | 0.0003 |  |  | 0.0162 | 0.0023 |
|  |  | B*45:01 | 0.1009 | 0.0491 | 0.0299 | 0.0628 | 0.0735 | 0.0077 | 0.0303 |
|  |  | B*51:01 | 0.0092 | 0.0173 | 0.0085 | 0.0152 | 0.0204 | 0.0527 | 0.0250 |
|  |  | B*53:01 | 0.0872 | 0.0636 | 0.0183 | 0.0974 | 0.0973 | 0.0096 | 0.0930 |
|  |  | B*58:02 | 0.0550 | 0.1127 | 0.1143 | 0.0801 | 0.0531 | 0.0006 | 0.0235 |

### Genetic Basis for Observed Differences

Furthermore, this study employed the Hardy-Weinberg equilibrium (HWE), Neutrality test of homozygosity, haplotypes and pairwise linkage disequilibrium test and inheritance patterns of alleles at different loci to gain an insight into the basis for the observed genetic diversity in each population at different loci. The HWE and Neutrality test of homozygosity are fundamental principles in population genetics and serve several important purposes in diversity studies [32, 33]. The HWE and Neutrality test were both performed at different HLA loci on the African populations. Significant deviations from expected HWE heterozygosity were observed in the distribution of genotypes of *HLA-C* in South Africa (Table 3) and the Ngoni tribe at HLA-B (Table 4).

**Table 3.** Exact test using Markov chain for HWE parameters for the five countries.

| Country | No of Genotype | Locus |  |  |  |  |  |  |  |  |
| --- | --- | --- | --- | --- | --- | --- | --- | --- | --- | --- |
|  |  | A |  |  | B |  |  | C |  |  |
|  |  | Obs. Het. | Exp. Het. | P-value (Adj) | Obs. Het. | Exp. Het. | P-value (Adj) | Obs. Het. | Exp. Het. | P-value (Adj) |
| Kenya | 109 | 0.9358 | 0.9299 | 0.8451 | 0.9633 | 0.9470 | 0.4505 | 0.9174 | 0.9143 | 0.6548 |
| Rwanda | 173 | 0.9249 | 0.9256 | 0.3050 | 0.9364 | 0.9459 | 0.8707 | 0.8786 | 0.8855 | 0.6548 |
| RSA | 1640 | 0.9348 | 0.9426 | 0.3750 | 0.9323 | 0.9355 | 0.4180 | 0.9012 | 0.9172 | 0.0155* |
| Uganda | 231 | 0.9351 | 0.9330 | 0.8324 | 0.9351 | 0.9499 | 0.7771 | 0.9048 | 0.9071 | 0.6548 |
| Zambia | 565 | 0.9221 | 0.9252 | 0.1445 | 0.9469 | 0.9424 | 0.4180 | 0.9062 | 0.9098 | 0.6390 |

\* Statistically significant. Obs. Het., observed heterozygosity; Exp. Het., expected heterozygosity.

**Table 4.**
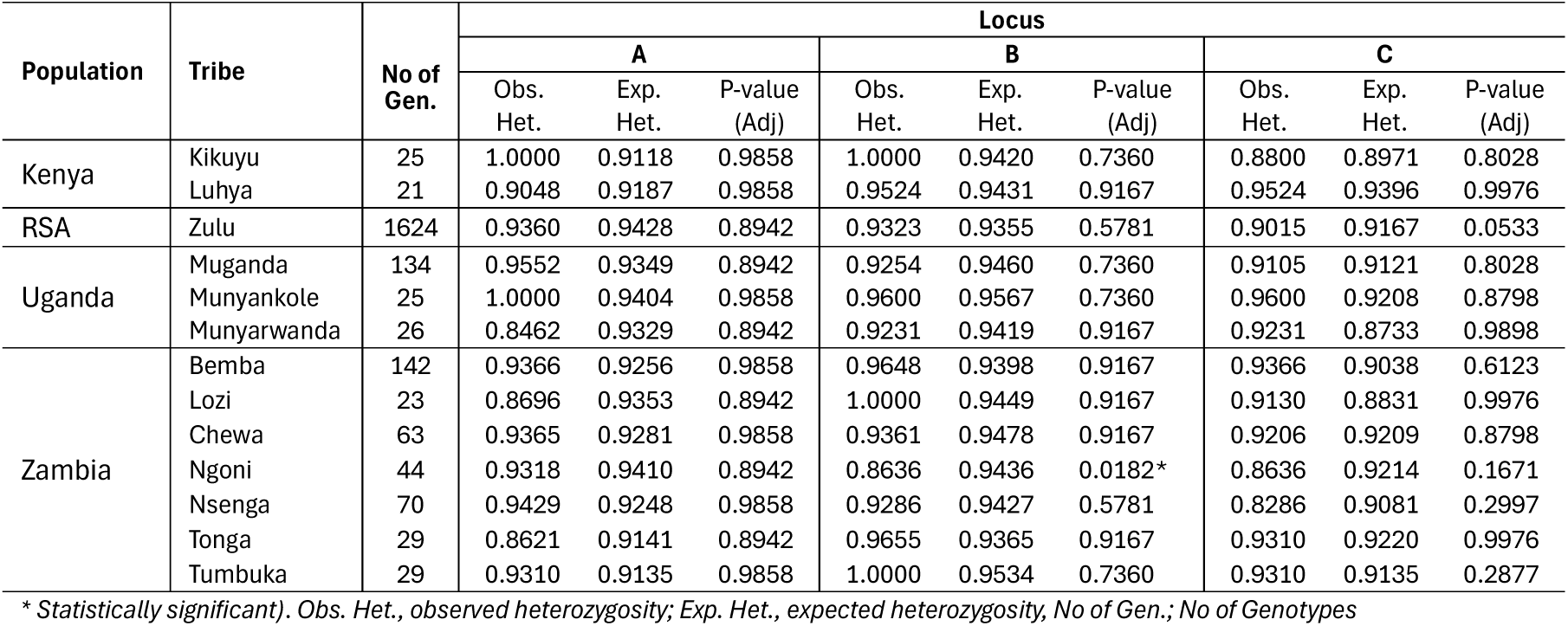
Exact test using Markov chain for HWE parameters for tribes.

| Population | Tribe | No of Gen. | Locus |  |  |  |  |  |  |  |  |
| --- | --- | --- | --- | --- | --- | --- | --- | --- | --- | --- | --- |
|  |  |  | A |  |  | B |  |  | C |  |  |
|  |  |  | Obs. Het. | Exp. Het. | P-value (Adj) | Obs. Het. | Exp. Het. | P-value (Adj) | Obs. Het. | Exp. Het. | P-value (Adj) |
| Kenya | Kikuyu | 25 | 1.0000 | 0.9118 | 0.9858 | 1.0000 | 0.9420 | 0.7360 | 0.8800 | 0.8971 | 0.8028 |
|  | Luhya | 21 | 0.9048 | 0.9187 | 0.9858 | 0.9524 | 0.9431 | 0.9167 | 0.9524 | 0.9396 | 0.9976 |
| RSA | Zulu | 1624 | 0.9360 | 0.9428 | 0.8942 | 0.9323 | 0.9355 | 0.5781 | 0.9015 | 0.9167 | 0.0533 |
| Uganda | Muganda | 134 | 0.9552 | 0.9349 | 0.8942 | 0.9254 | 0.9460 | 0.7360 | 0.9105 | 0.9121 | 0.8028 |
|  | Munyankole | 25 | 1.0000 | 0.9404 | 0.9858 | 0.9600 | 0.9567 | 0.7360 | 0.9600 | 0.9208 | 0.8798 |
|  | Munyarwanda | 26 | 0.8462 | 0.9329 | 0.8942 | 0.9231 | 0.9419 | 0.9167 | 0.9231 | 0.8733 | 0.9898 |
| Zambia | Bemba | 142 | 0.9366 | 0.9256 | 0.9858 | 0.9648 | 0.9398 | 0.9167 | 0.9366 | 0.9038 | 0.6123 |
|  | Lozi | 23 | 0.8696 | 0.9353 | 0.8942 | 1.0000 | 0.9449 | 0.9167 | 0.9130 | 0.8831 | 0.9976 |
|  | Chewa | 63 | 0.9365 | 0.9281 | 0.9858 | 0.9361 | 0.9478 | 0.9167 | 0.9206 | 0.9209 | 0.8798 |
|  | Ngoni | 44 | 0.9318 | 0.9410 | 0.8942 | 0.8636 | 0.9436 | 0.0182* | 0.8636 | 0.9214 | 0.1671 |
|  | Nsenga | 70 | 0.9429 | 0.9248 | 0.9858 | 0.9286 | 0.9427 | 0.5781 | 0.8286 | 0.9081 | 0.2997 |
|  | Tonga | 29 | 0.8621 | 0.9141 | 0.8942 | 0.9655 | 0.9365 | 0.9167 | 0.9310 | 0.9220 | 0.9976 |
|  | Tumbuka | 29 | 0.9310 | 0.9135 | 0.9858 | 1.0000 | 0.9534 | 0.7360 | 0.9310 | 0.9135 | 0.2877 |

\* Statistically significant). Obs. Het., observed heterozygosity; Exp. Het., expected heterozygosity, No of Gen.; No of Genotypes

Also, the Neutrality test of homozygosity showed significant deviations from expected homozygosity in Kenya, South Africa and Uganda at locus A and in Rwanda and Uganda at locus B (Table 5). At tribal level, similar deviations were also observed at locus B for Muganda, Nsenga and Tumbuka (Table 6). Also, Chewa and Muganda tribes had significant deviations from expected homozygosity at locus C (Table 6).

**Table 5.** Slatkin’s implementation of EW homozygosity test of neutrality for the five African countries.

| Country | Locus |  |  |  |  |  |  |  |  |  |  |  |
| --- | --- | --- | --- | --- | --- | --- | --- | --- | --- | --- | --- | --- |
|  | A |  |  |  | B |  |  |  | C |  |  |  |
|  | Obs. (Homo) F | Exp. (Homo) F | Fnd | p-value (Adj) | Obs. (Homo) F | Exp. (Homo) F | Fnd | p-value (Adj) | Obs. (Homo) F | Exp. (Homo) F | Fnd | p-value (Adj) |
| Kenya | 0.0744 | 0.1238 | -1.2527 | 0.0430* | 0.0574 | 0.0764 | -0.9347 | 0.1231 | 0.0899 | 0.1431 | -1.1121 | 0.0591 |
| Rwanda | 0.0771 | 0.1060 | -0.8577 | 0.1502 | 0.0568 | 0.0955 | -1.3298 | 0.0338* | 0.1171 | 0.1703 | -0.8657 | 0.1440 |
| RSA | 0.0577 | 0.1111 | -1.4135 | 0.0240* | 0.0648 | 0.0940 | -0.9500 | 0.1231 | 0.0831 | 0.1492 | -1.1969 | 0.0503 |
| Uganda | 0.0690 | 0.1147 | -1.2026 | 0.0430* | 0.0522 | 0.0861 | -1.3086 | 0.0338* | 0.0948 | 0.1816 | -1.2900 | 0.0503 |
| Zambia | 0.0756 | 0.1204 | -1.0813 | 0.0776 | 0.0584 | 0.0945 | -1.1905 | 0.0583 | 0.0910 | 0.1613 | -1.1671 | 0.0503 |

\* Statistically significant.

**Table 6.**
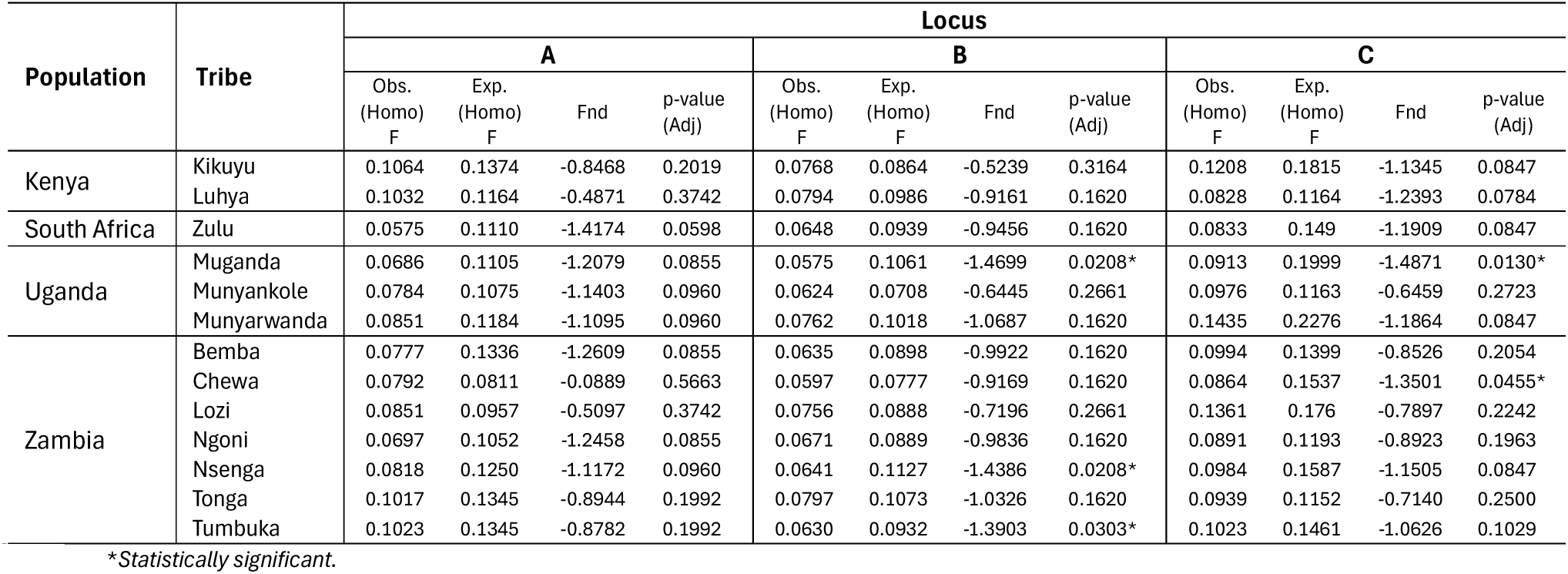
Slatkin’s implementation of EW homozygosity test of neutrality for tribes.

Haplotypes and linkage disequilibrium analysis helps in understanding the genetic variation of alleles that are inherited together on the same chromosome and non-random associations among alleles at different loci respectively. The haplotypic associations of the HLA class I region were also investigated. While the full list of haplotypes is detailed in the **Supplementary Tables 4, 5, 6,and 7**, topmost estimated two and three loci haplotypes in each population are summarized in Table 7. At loci A:B, A:C and A:B:C, haplotypes *A*30:01∼B*42:01, A*30:01∼C*17:01* and *A*30:01∼B*42:01∼C*17:01* were the topmost in the South Africa and Zambia populations. Similarly, Haplotypes *A*02:01∼B*15:03* and *A*02:01∼B*15:03∼C*02:10* were detected at similar frequencies as topmost haplotypes in the Rwanda and Uganda populations. South Africa and Rwanda reported the same top haplotypes at locus pair B:C (*B*58:02∼C*06:02*) at similar frequencies. All the populations reported different topmost haplotypes at three loci association (Table 7). Between the two loci, the strongest estimated associations were those between alleles of HLA-B and C (Table 8).

**Table 7.** Topmost haplotypes at different loci across populations.

| Populations | Loci |  |  |  |  |  |  |  |
| --- | --- | --- | --- | --- | --- | --- | --- | --- |
|  | A:B |  | A:C |  | B:C |  | A:B:C |  |
|  | A~B | HF | A~C | HF | B~C | HF | A~B~C | HF |
| Kenya | A*68:02~B*15:10 | 0.0548 | A*68:02~C*03:04 | 0.0596 | B*42:01~C*17:01 | 0.0734 | A*68:02~B*27:03~C*02:02 | 0.0596 |
| Rwanda | A*02:01~B*15:03 | 0.0513 | A*02:02~C*06:02 | 0.0571 | B*58:02~C*06:02 | 0.1098 | A*02:01~B*15:03~C*02:10 | 0.0484 |
| RSA | A*30:01~B*42:01 | 0.0465 | A*30:01~C*17:01 | 0.0355 | B*58:02~C*06:02 | 0.1134 | A*30:01~B*42:01~C*17:01 | 0.0275 |
| Uganda | A*02:01~B*15:03 | 0.0426 | A*02:01~C*02:10 | 0.0388 | B*53:01~C*04:01 | 0.0824 | A*02:01~B*15:03~C*02:10 | 0.0405 |
| Zambia | A*30:01~B*42:01 | 0.0480 | A*30:01~C*17:01 | 0.0557 | B*15:03~C*02:10 | 0.0937 | A*30:01~B*42:01~C*17:01 | 0.0463 |
| Caucasian | A*01:01~B*08:01 | 0.0501 | A*01:01~C*07:01 | 0.0587 | B*07:02~C*07:02 | 0.0929 | A*01:01~B*08:01~C*07:01 | 0.0503 |
| AFAM | A*30:02~B*57:03 | 0.0267 | A*02:01~C*16:01 | 0.0234 | B*53:01~C*04:01 | 0.0763 | A*33:03~B*53:01~C*04:01 | 0.0151 |

HF: Haplotype Frequency

**Table 8.** Pairwise linkage disequilibrium across countries.

| Country | Loci |  |  |  |  |  |  |  |  |
| --- | --- | --- | --- | --- | --- | --- | --- | --- | --- |
|  | A:B |  |  | A:C |  |  | B:C |  |  |
| | $D'$ | $W_n$ | P-value (Adj) | $D'$ | $W_n$ | P-value (Adj) | $D'$ | $W_n$ | P-value (Adj) |
| Kenya | 0.7743 | 0.5234 | < 0.001 | 0.6870 | 0.4210 | < 0.001 | 0.8926 | 0.8112 | < 0.001 |
| Rwanda | 0.6618 | 0.4513 | < 0.001 | 0.5971 | 0.4938 | < 0.001 | 0.9094 | 0.7817 | < 0.001 |
| South Africa | 0.6100 | 0.3850 | < 0.001 | 0.5597 | 0.3884 | < 0.001 | 0.8819 | 0.5934 | < 0.001 |
| Uganda | 0.6732 | 0.4232 | < 0.001 | 0.6024 | 0.3915 | < 0.001 | 0.8987 | 0.7488 | < 0.001 |
| Zambia | 0.5638 | 0.4275 | < 0.001 | 0.5325 | 0.3198 | < 0.001 | 0.8748 | 0.6897 | < 0.001 |

Pairwise linkage disequilibrium measured by Hedrick’s and Crammer’s statistics at all loci across populations were all statistically significant (*P* < 0.001) as presented in Table 8. Few loci such as A:B in Lozi, Tonga and Tumbuka, A:C in Tonga, show random association (not significant) between alleles (Table 9).

**Table 9.** Pairwise linkage disequilibrium across tribes.

| Population | Tribe | Loci |  |  |  |  |  |  |  |  |
| --- | --- | --- | --- | --- | --- | --- | --- | --- | --- | --- |
|  |  | A:B |  |  | A:C |  |  | B:C |  |  |
| | | $D'$ | $W_n$ | P-value (adj) | $D'$ | $W_n$ | P-value (adj) | $D'$ | $W_n$ | P-value (adj) |
| Kenya | Kikuyu | 0.8714 | 0.7535 | 0.0065 | 0.7938 | 0.6163 | 0.0498 | 0.9734 | 0.8947 | <0.001 |
|  | Luhya | 0.9431 | 0.7011 | <0.001 | 0.9522 | 0.6504 | 0.0065 | 0.9635 | 0.9014 | <0.001 |
| RSA | Zulu | 0.6117 | 0.4061 | <0.001 | 0.5624 | 0.3899 | <0.001 | 0.8831 | 0.6012 | <0.001 |
| Uganda | Muganda | 0.7240 | 0.4205 | <0.001 | 0.6697 | 0.5050 | <0.001 | 0.8994 | 0.8154 | <0.001 |
|  | Munyankole | 0.9280 | 0.7406 | 0.0231 | 0.8924 | 0.6611 | <0.001 | 0.9808 | 0.8862 | <0.001 |
|  | Munyarwanda | 0.8520 | 0.6791 | 0.0312 | 0.7936 | 0.6764 | 0.0169 | 0.9431 | 0.8774 | <0.001 |
| Zambia | Bemba | 0.6978 | 0.4596 | <0.001 | 0.6295 | 0.3748 | <0.001 | 0.8891 | 0.7498 | <0.001 |
|  | Chewa | 0.8346 | 0.6363 | <0.001 | 0.7634 | 0.6442 | <0.001 | 0.9733 | 0.8984 | <0.001 |
|  | Lozi | 0.9131 | 0.6916 | 0.1712* | 0.8173 | 0.7436 | 0.0425 | 0.9237 | 0.8416 | <0.001 |
|  | Ngoni | 0.8305 | 0.6637 | <0.001 | 0.8007 | 0.6433 | 0.0037 | 0.9506 | 0.8323 | <0.001 |
|  | Nsenga | 0.7514 | 0.5362 | <0.001 | 0.6890 | 0.4343 | <0.001 | 0.9114 | 0.7371 | <0.001 |
|  | Tonga | 0.8120 | 0.6539 | 0.1474* | 0.7742 | 0.5671 | 0.4344* | 0.9189 | 0.7068 | <0.001 |
|  | Tumbuka | 0.8561 | 0.6335 | 0.1334* | 0.8084 | 0.6253 | 0.0087 | 0.9204 | 0.7830 | <0.001 |

\*Statistically not significant

Additionally, this study investigated alleles that are unique to different populations (see Supplementary Tables 8 and 9). Based on the sample sizes of each population in this study, it was observed that certain alleles were unique to different populations.

Furthermore, Alluvial plots were employed to present the inheritance patterns of alleles observed in this study. Each block size in the alluvial plot represents the frequency of the corresponding alleles and the thickness of the flow streams denotes the frequency of alleles inheritance pattern. These provide an understanding of predicting the likelihood of inheriting specific traits or conditions [34, 35]. Figure 5 presents inheritance patterns of HLA-B alleles as observed in the African populations (**see Supplementary Figures 3, 4, and 5 for full list**). In Kenya, it was observed that *HLA-B*07:02* was inherited more often with *HLA-B*45:01* than other alleles. Also, *HLA-B*45:01* (Allele_1 and Allele_2) was observed to be inherited more often with *HLA-B*15:10*, and *HLA-B*58:02*. Furthermore, *HLA-B*42:01* was observed to have more inheritance patterns with *HLA-B*53:01* and *HLA-B*58:01* alleles. Similarly, *HLA-B*15:03* and *HLA-B*58:02* alleles were observed to be inherited together with most of the alleles in Rwanda. In Ugandan and Zambian populations, *HLA-B*53:01* had the highest inheritance pattern with other alleles. *HLA-B*58:02* had the highest pattern of inheritance in both Rwanda and South Africa followed by *HLA-B*15:03* and *HLA-B*42:01* in Rwanda and South Africa respectively.

**Figure 5.**
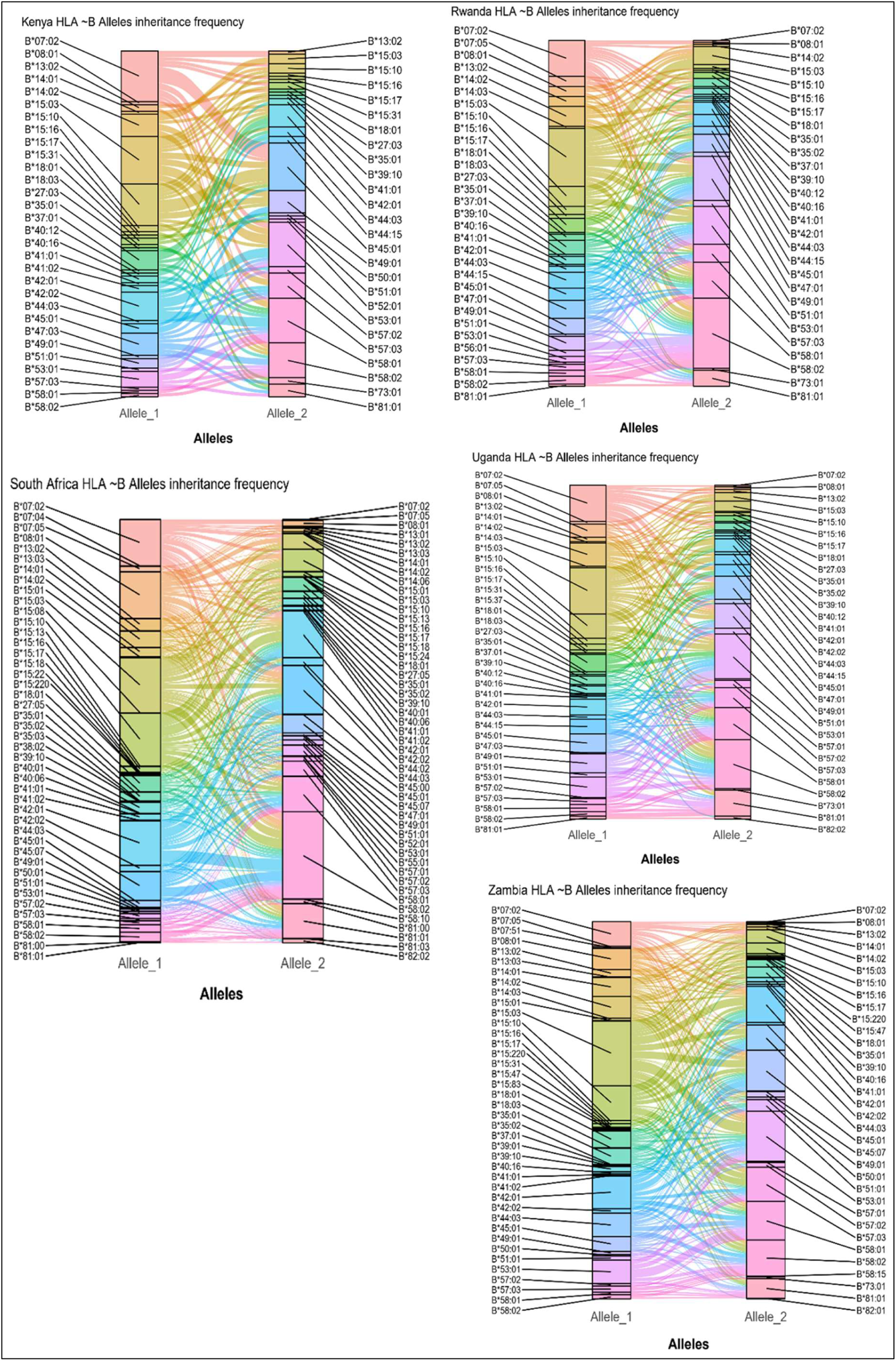
plots showing frequency how HLA ∼B alleles were inherited together by participants in each country.

## DISCUSSION

This study adopted several population genetic diversity approaches to investigate class I HLA diversity in the eastern and southern African populations compared to the US populations.

This study observed differences in allele frequencies in all populations. Distribution of allele frequencies are influenced by several factors such as genetic drift, gene flow, mutation, population history, natural selection, making each population genetically unique [36–38]. Allele frequencies vary across populations and the topmost 5% frequent alleles reported in this study have also been reported in other studies [19, 39–41] at higher or lower frequencies. For example, *HLA-A*02:01* is a common allele of the *HLA-A* gene, playing a crucial role in the immune system [42, 43]. The high prevalence of *HLA-A*02:01* in a population has been linked to higher risk of certain cancers [44]. This distinction among the alleles frequencies across populations is a testament to the HLA genetic architectural diversity among the populations.

The Jaccard index heatmap shows various levels of allelic similarities among populations (regions, countries and tribes). The heatmap indicated that the Caucasian and AFAM are dissimilar to the African population due to extremely low similarity indices observed at all loci. This affirms the allelic diversity between the African and US populations and suggests that there are several uncommon alleles between the two populations. Population comparisons based on haplotype frequencies using MDS showed distinct genetic differences both within African populations and between African and US populations. The Cartograph clearly shows the distinction between the African and United State populations. The Caucasian population show high genetic distances to the African populations at all loci which indicated high diversity between the two populations. The African American population though genetically close to

African population due to their historical background [45], still maintain a level of distinction which suggests a non-representative of the African populations.

The Shannon and Simpson indices affirm the polymorphic status of each HLA locus and suggested different levels of diversity within each population. Interestingly, despite the lower number of HLA-A alleles detected in the South African populations, the Shannon index shows that this locus displayed relatively evened allele distributions which resulted in higher diversity than *HLA-B* that had more alleles.

The highest values of the Jaccard index in the African populations were observed among the eastern African region at all loci. This suggests high similarity in terms of their combination of common alleles. Majority of the southern African countries had low similarity due to uncommon alleles between them. At the tribal level, the Zulu tribe also exhibited low similarity to other tribes within the African region but maintained relatively high similarities with the Zambian tribes at all loci. This affirms the closeness between the two populations at both country and tribal levels. Summarily, the Jaccard indices observed at the tribal levels also affirm the existence of allelic diversity among tribes of the same countries within African populations. The genetic distances observed in the cartograph suggest allelic diversity among the populations as previously established by other analyses in this study. High diversity was observed in some HLA loci (A:B and A:C) than others (B:C and A:B:C) among African countries. Countries from the same region tended to be in the same location on the cartograph for haplotypes A∼B and B∼C. This suggests similar genetic diversities between those countries in the same region. Also, South Africa seemed to have a close genetic distance to Zambia at loci A:B and A:C compared to other countries and a closer genetic distance at loci A:B:C. Interestingly, there was a wide genetic distance between the two countries at loci B:C. This could be linked to some allelic bias towards

South Africa compared to Zambia even though both countries are from the same African region. Kenya showed closer genetic distance to Zambia at loci A:B and A:C compared to any other African countries. Similar closeness was also observed at loci B:C between Kenya and Uganda. This suggests low diversity between the two countries at those loci. Furthermore, there was high diversity between Kenya and all other African populations at loci A:B:C. Also, we observed distinct levels of diversity between Rwanda and Uganda at different loci. While high diversity was observed at loci A:C, relatively low diversity was observed between the two countries at loci A:B, B:C and A:B:C.

Additionally, the rarefaction curves support the comparison of allelic diversity between populations (countries and tribes). Although, by observing the shape of the curves, we can infer that allelic variants have been observed within a given number of samples, yet more participants are required to the HLA typed to observed the unique alleles in all the African populations. Furthermore, diversity at locus B is more observed across populations due to the polymorphic nature of alleles at that locus [46].

Generally, the absence of Bw4 and Bw6 alleles in African populations indicates non-expression of serological markers at the respective locus, which can affect organ transplantation compatibility, immune responses and disease susceptibility within the continent [47, 48]. Similarly, the absence of *HLA-B*27:05* in African is supported by the uncommon presentation of ankylosing spondylitis (AS) disease [49]. *HLA-B*27:05* has been reported in literature to be associated with AS [50, 51] and high prevalence of AS disease in Caucasian and African American populations is said to be associated with *HLA-B*27:05* [52, 53].

Also, the study observed significant deviation from Hardy-Weinberg equilibrium in South Africa population at locus C. Similar deviations were observed in Ngoni at locus B. Potential causes of significant deviation from Hardy-Weinberg equilibrium have been mentioned in literature [19, 40, 41]. Deviations from HWE at these loci in the two populations might indicate inbreeding, which can reduce genetic diversity and the population’s ability to adapt to environmental changes at these loci [54, 55]. However, due to the retrospective nature of this study, we acknowledge allelic bias and/or HLA genotyping error as major potential causes of the deviations as also reported in literatures [56]. Ewens-Watterson Neutrality test of homozygosity was significant for different populations at different loci. The significant deviations observed for the different populations at different loci suggest balancing selection which helps in preserving multiple alleles at each locus, contributing to genetic diversity [57–60]. This is vital for the adaptability and long-term survival of populations, enabling them to cope with changing environments and disease pressure that are associated with alleles in that locus [61].

Also, top haplotypes observed between populations affirm the closeness among such populations at the respective locus. There was a strong LD between all the locus pair across populations in this study except for Lozi, Tonga, and Tumbuka at locus A:B and Tonga at locus A:C. It is reported that haplotype frequencies are influenced by allele frequencies, LD, samples sizes, completeness of HLA data etc. [62–64]. The results show genetic variants in high non-random associations being less likely to be separated by a recombination event and thus alleles of the variants are more commonly inherited together than expected [65, 66]. Hedrick’s D′ weights alleles in each haplotype and Cramer’s V Statistic is a multi-allelic correlation measure between pairs of loci [41]. Also, haplotype diversity coupled with highly significant LD might provide insight into Negative (or purifying) selection in the HLA genomic region [67]. This could also be linked to background selection where linked allelic variations are lost during negative selection process [68]. Similar pattern of results was observed at the tribal level which also indicates genetic diversity among tribes at all loci.

The discrepancies in the unique alleles observed in the groups of population might be due to sample sizes of the populations in this study. Hence, larger sample sizes with more African countries need to be studied to get a comprehensive picture of HLA genetic diversity across Africa

This study only looked at classical Class I *HLA* genes and the patterns of alleles inheritance at each locus observed in this study needs to be studied in more details. Also, More researches need to look at non-classical genes and Class II genes as it will help in unravelling the genetic profiles in terms of disease susceptibility and protection in each population. This will assist in understanding genetic diversity and inform in *HLA* population-based therapeutic development for each country.

### Limitations of study

This study had limitation in terms of samples sizes in the tribal populations which could have increase the understanding of the diversity among enough tribes within each country. Hence, large sample sizes of HLA data at tribal level are needed to fully understand their respective diversity. Additionally, the imbalance sample sizes among populations might have influenced the number alleles, alleles and haplotype frequencies within each population. However, the limitations observed do not affect the importance of understanding HLA diversity in the African subregion as presented in this study.

## Conclusion

In this study, we have established HLA diversity in the Eastern and Southern African region of the African continent. Comparison of the HLA data at both country and tribal levels suggest genetic differences within the African populations and uniqueness of the Eastern and Southern African populations relative to the US-based African populations. These analyses demonstrate the limitations of applying HLA data from one region to another, reinforcing the necessity of collecting high-quality HLA data from all regions of Africa and its varied ethnicities. Comprehensive data collection is crucial for enhancing vaccine design and advancing our understanding of HLA disease associations, ultimately improving healthcare outcomes across the continent. Finally, due to genetic admixture, cautions must be made against extrapolating HLA data from other continents to inform African vaccine development.

## MATERIALS AND METHODS

### Population and Sample

The Class I HLA data used in this study were obtained from a preliminary study of our HLA typing project and also from our collaborators across five distinct cohorts within African populations and two ethnic groups in the United States, all of which are part of HIV research cohorts. The African cohorts comprise Centre for The Aids Programme of Research In South Africa (CAPRISA), International AIDS Vaccine Initiative (IAVI), Female Rising through Education, Support and Health (FRESH), and Sinikithemba in South Africa. The ethnic groups from the US are the African Americans (AFAM) and Caucasians (CAU) [20]. Necessary approvals were granted for all the HLA studies across the different cohorts. The present study includes 2,718 anonymous samples from apparent unrelated subjects across the different cohorts. African samples were obtained from three eastern and two southern African countries and are distributed as follows; Kenya (*n* = 106), Rwanda (*n* = 173), Uganda (*n* = 231), South Africa – RSA (*n* = 1640) and Zambia (*n* = 565). Of the five countries sampled within the African sub-region, tribal information was obtained from four countries excluding Rwanda due to historical development. The ethnic groups sampled within the four countries are Bemba, Chewa, Kikuyu, Lozi, Luhya, Muganda, Munyankole, Munyarwanda, Ngoni, Nsenga, Tonga, Tumbuka and Zulu. Similarly, the US ethnic groups were distributed as CAU (*n* = 1765) and AFAM (*n* = 661). In accordance to the World Medical Association Declaration of Helsinki [69], participants’ personal identifiers were not accessed to maintain confidentiality.

### Data Cleaning and Validation

The HLA data used in this study was examined for inconsistencies and an *in silico* method (expert knowledge) [70] was used to resolve the ambiguities encountered. Few samples were duplicated with similar allelic information and participants with more allelic information were retained for the study. Otherwise, only one sample was retained in the case of same allelic information in the sample. Also, duplicate samples with different allelic information and samples with partially or entirely missing allelic information were excluded from the analysis. Furthermore, the HLA data was analysed at 4-digit resolution in this study.

All the HLA data used in this study were checked for allele validity, and all allele nomenclature reported prior to 2010 were updated using current nomenclature conversion tables and conversion tools provided by IMGT/HLA databased (IMGT/HLA Database, IPD-IMGT/HLA 3.56, release of January 2024, https://www.ebi.ac.uk/ipd/imgt/hla/alleles/). Similarly, haplotype nomenclature was done in accordance with the 2013 report [71] aimed at organizing and discriminating phased genes, genotypes, and ambiguous assignments.

### Statistical Analysis

Allele frequencies were estimated by direct counting using Python for population genomics (PyPop) version 1.0.0 [72]. The haplotypes and haplotype frequencies (HF) were estimated by resolving phase and allelic ambiguities using the expectation-maximization (EM) steps with progressive insertion algorithm by setting the posterior probability to 0.0001 in the haplo.stats version 1.9.5.1 R package [73]. The HLA data were converted to Arlequin version 3.5.2 software [74] input files using CREATE software version 1.37 [75] to examine deviations from Hardy-Weinberg equilibrium (HWE) adopting a modification of the Markov random walk algorithm with 100 000 dememorization steps [76]. Estimation of relative delta (*D*′) and Cramer’s V Statistic (*W*_n_) values to measure pairwise linkage disequilibrium (LD) between pairs of alleles of different loci and their statistical significance were calculated using Hedrick’s [77] and Cramer’s [78] estimators as previously described in literature [39, 79]. The Ewen-Watterson neutrality test of homozygosity was implemented in PyPop using the Slatkin principle of implementation [80, 81]. Multiple comparisons of both LD and Neutrality tests of homozygosity were both addressed via Benjamini & Hochberg correction method [82]. Aplha diversity indices such as specie richness - number of alleles [83, 84], Shannon index – entropy [85], and Simpson **(Gini-Simpson)** index - probability that two alleles taken from the sample at random are of different types [86, 87] were all used to measure within population diversities. The Jaccard similarity index [88], a measure of beta diversity, was employed to determine heterogeneity between the populations. Furthermore, a rarefaction analysis to gain quantitative insights into the number of alleles that were observed in each population as a function of the number of participants was also determined. Similarly, as a measure of genetic distance between populations, haplotype frequency data from each country were dimensionality reduced using classical multidimensional scaling (MDS) to create a 2-dimensional genetic cartograph. Based on the analysis, two countries are close to each other on the map if the distribution of the HLA alleles in these two countries are close to each other, relative to the distribution observed in the other countries.

## ACKNOWLEDGEMENTS

We would like to express our sincere gratitude to Dr. Mary Carrington from the HLA Immunogenetics Section in the Laboratory of Integrative Cancer Immunology at the National Cancer Institute (NCI) for providing the Caucasian and African American HLA data, as well as for her invaluable technical support. We also thank Dr. Bruce Walker from the Ragon Institute of Massachusetts General Hospital, Massachusetts Institute of Technology, for supplying the HLA data from the elite controllers and the FRESH study cohort. Our appreciation extends to Dr. Thumbi Ndung’u of the Africa Health Research Institute (AHRI) in Durban, South Africa, for providing the Sinikithemba HLA data. Additionally, we are grateful to the International AIDS Vaccine Initiative (IAVI) and the Centre for the AIDS Programme of Research in South Africa (CAPRISA) for contributing some of the HLA data utilized in this study.

The authors would like to acknowledge the following funding sources that supported the research, authorship, and publication of this article: The Bill and Melinda Gates Foundation (Grant #INV-048833 to ZMN; Grant #INV-027090 to ZMN; Grant #INV-050722 to ZMN). National Institutes of Health (NIH/NIAID)Grant # [R01AI181690] to ZMN: Grant #R01A1145305 to ZMN), Sub-Saharan African Network for TB/HIV Research Excellence (SANTHE), Collaborative award (grant # SANTHE COL018).

## AUTHOR CONTRIBUTIONS

ZMN conceptualized the study design and secured funding. ZMN, VR, OH, and TN initiated the study design, with TN coordinating the study. ZMN and VR obtained data from collaborators. ZMN, TN, and NN performed the HLA typing, while AWB and OH analysed the data. TN, NN, and AWB drafted the original manuscript, with ZMN supervising the writing process. VR, TN, NN, and AWB reviewed and edited the original manuscript.

## Abbreviations

RSA: Republic of South Africa
HF: Haplotype Frequency
CAU: Caucasian
AFAM: African American
LD: Linkage Disequilibrium

## REFERENCES

1. Ben Bnina A, Yessine A, El Bahri Y, Chouchene S, Ben Lazrek N, Mimouna M, et al. Contribution of HLA class I (A, B, C) and HLA class II (DRB1, DQA1, DQB1) alleles and haplotypes in exploring ethnic origin of central Tunisians. BMC Med Genomics. 2024;17(1):65.

2. Arrieta-Bolanos E, Hernandez-Zaragoza DI, Barquera R. An HLA map of the world: A comparison of HLA frequencies in 200 worldwide populations reveals diverse patterns for class I and class II. Front Genet. 2023;14:866407.

3. Tattersall I. Exploring the “Cradle of Humankind”. Evolution: Education and Outreach. 2010;3(3):466–7.

4. Campbell MC, Tishkoff SA. The evolution of human genetic and phenotypic variation in Africa. Curr Biol. 2010;20(4):R166–73.

5. Campbell MC, Tishkoff SA. African genetic diversity: implications for human demographic history, modern human origins, and complex disease mapping. Annu Rev Genomics Hum Genet. 2008;9:403–33.

6. Cherry M. Human evolution: The cradle of humankind revisited. Nature. 2015;523(7558):33-.

7. Gonzalez-Galarza FF, McCabe A, Santos E, Jones J, Takeshita L, Ortega-Rivera ND, et al. Allele frequency net database (AFND) 2020 update: gold-standard data classification, open access genotype data and new query tools. Nucleic Acids Res. 2020;48(D1):D783–d8.

8. Gonzalez-Galarza FF, McCabe A, Melo Dos Santos EJ, Jones AR, Middleton D. A snapshot of human leukocyte antigen (HLA) diversity using data from the Allele Frequency Net Database. Hum Immunol. 2021;82(7):496–504.

9. Janse van Rensburg WJ, de Kock A, Bester C, Kloppers JF. HLA major allele group frequencies in a diverse population of the Free State Province, South Africa. Heliyon. 2021;7(4):e06850.

10. Ali AA, Aalto M, Jonasson J, Osman A. Genome-wide analyses disclose the distinctive HLA architecture and the pharmacogenetic landscape of the Somali population. Sci Rep. 2020;10(1):5652.

11. Blackwell JM, Jamieson SE, Burgner D. HLA and infectious diseases. Clin Microbiol Rev. 2009;22(2):370-85, Table of Contents.

12. Bodis G, Toth V, Schwarting A. Role of Human Leukocyte Antigens (HLA) in Autoimmune Diseases. Rheumatol Ther. 2018;5(1):5–20.

13. Ghattaoraya GS, Middleton D, Santos EJ, Dickson R, Jones AR, Alfirevic A. Human leucocyte antigen-adverse drug reaction associations: from a perspective of ethnicity. Int J Immunogenet. 2017;44(1):7–26.

14. Creary LE, Sacchi N, Mazzocco M, Morris GP, Montero-Martin G, Chong W, et al. High-resolution HLA allele and haplotype frequencies in several unrelated populations determined by next generation sequencing: 17th International HLA and Immunogenetics Workshop joint report. Hum Immunol. 2021;82(7):505–22.

15. Goulder PJ, Walker BD. HIV and HLA class I: an evolving relationship. Immunity. 2012;37(3):426–40.

16. Darbas S, Inan D, Kilinc Y, Arslan HS, Ucar F, Boylubay O, et al. Relationship of HLA- B alleles on susceptibility to and protection from HIV infection in Turkish population. North Clin Istanb. 2023;10(1):67–73.

17. Kiepiela P, Leslie AJ, Honeyborne I, Ramduth D, Thobakgale C, Chetty S, et al. Dominant influence of HLA-B in mediating the potential co-evolution of HIV and HLA. Nature. 2004;432(7018):769-75.

18. Leslie A, Matthews PC, Listgarten J, Carlson JM, Kadie C, Ndung’u T, et al. Additive contribution of HLA class I alleles in the immune control of HIV-1 infection. J Virol. 2010;84(19):9879–88.

19. Cao K, Hollenbach J, Shi X, Shi W, Chopek M, Fernández-Viña MA. Analysis of the frequencies of HLA-A, B, and C alleles and haplotypes in the five major ethnic groups of the United States reveals high levels of diversity in these loci and contrasting distribution patterns in these populations. Hum Immunol. 2001;62(9):1009–30.

20. Pereyra F, Jia X, McLaren PJ, Telenti A, de Bakker PI, Walker BD, et al. The major genetic determinants of HIV-1 control affect HLA class I peptide presentation. Science. 2010;330(6010):1551-7.

21. GRAGERT L, DiPrima S, Albrecht M, Maiers M, Kalaycio M, Hill BT. HLA Is a Determinant Of The Ethnic Predisposition Of Chronic Lymphocytic Leukemia. Blood. 2013;122(21):1620-.

22. Laland KN, Uller T, Feldman MW, Sterelny K, Müller GB, Moczek A, et al. The extended evolutionary synthesis: its structure, assumptions and predictions. Proc Biol Sci. 2015;282(1813):20151019.

23. Scott-Phillips TC, Laland KN, Shuker DM, Dickins TE, West SA. The niche construction perspective: a critical appraisal. Evolution. 2014;68(5):1231–43.

24. Sidney J, Peters B, Frahm N, Brander C, Sette A. HLA class I supertypes: a revised and updated classification. BMC Immunology. 2008;9(1):1.

25. Vinkšel M, Writzl K, Maver A, Peterlin B. Improving diagnostics of rare genetic diseases with NGS approaches. J Community Genet. 2021;12(2):247–56.

26. Strianese O, Rizzo F, Ciccarelli M, Galasso G, D’Agostino Y, Salvati A, et al. Precision and Personalized Medicine: How Genomic Approach Improves the Management of Cardiovascular and Neurodegenerative Disease. Genes (Basel). 2020;11(7).

27. Hurley CK. Naming HLA diversity: A review of HLA nomenclature. Human Immunology. 2021;82(7):457–65.

28. Lutz CT. Human leukocyte antigen Bw4 and Bw6 epitopes recognized by antibodies and natural killer cells. Curr Opin Organ Transplant. 2014;19(4):436–41.

29. Goulder Philip JR, Walker Bruce D. HIV and HLA Class I: An Evolving Relationship. Immunity. 2012;37(3):426–40.

30. Lutz CT. Human leukocyte antigen Bw4 and Bw6 epitopes recognized by antibodies and natural killer cells. Current Opinion in Organ Transplantation. 2014;19(4):436–41.

31. Valenzuela-Ponce H, Alva-Hernández S, Garrido-Rodríguez D, Soto-Nava M, García-Téllez T, Escamilla-Gómez T, et al. Novel HLA class I associations with HIV-1 control in a unique genetically admixed population. Sci Rep. 2018;8(1):6111.

32. Wigginton JE, Cutler DJ, Abecasis GR. A note on exact tests of Hardy-Weinberg equilibrium. Am J Hum Genet. 2005;76(5):887–93.

33. Nielsen R. Statistical tests of selective neutrality in the age of genomics. Heredity. 2001;86(6):641–7.

34. Woodward AA, Urbanowicz RJ, Naj AC, Moore JH. Genetic heterogeneity: Challenges, impacts, and methods through an associative lens. Genet Epidemiol. 2022;46(8):555–71.

35. Genetic A, The New England Public Health Genetics Education C. Genetic Alliance Monographs and Guides. Understanding Genetics: A New England Guide for Patients and Health Professionals. Washington (DC): Genetic Alliance Copyright © 2008, Genetic Alliance.; 2010.

36. Sork VL. Gene flow and natural selection shape spatial patterns of genes in tree populations: implications for evolutionary processes and applications. Evol Appl. 2016;9(1):291–310.

37. Mattei J, Parnell LD, Lai C-Q, Garcia-Bailo B, Adiconis X, Shen J, et al. Disparities in allele frequencies and population differentiation for 101 disease-associated single nucleotide polymorphisms between Puerto Ricans and non-Hispanic whites. BMC Genetics. 2009;10(1):45.

38. Marth GT, Czabarka E, Murvai J, Sherry ST. The Allele Frequency Spectrum in Genome-Wide Human Variation Data Reveals Signals of Differential Demographic History in Three Large World Populations. Genetics. 2004;166(1):351–72.

39. Bugawan TL, Klitz W, Blair A, Erlich HA. High-resolution HLA class I typing in the CEPH families: analysis of linkage disequilibrium among HLA loci. Tissue Antigens. 2000;56(5):392–404.

40. Tokić S, Žižkova V, Štefanić M, Glavaš-Obrovac L, Marczi S, Samardžija M, et al. HLA-A, -B, -C, -DRB1, -DQA1, and -DQB1 allele and haplotype frequencies defined by next generation sequencing in a population of East Croatia blood donors. Scientific Reports. 2020;10(1):5513.

41. Tshabalala M, Mellet J, Vather K, Nelson D, Mohamed F, Christoffels A, et al. High Resolution HLA ∼A, ∼B, ∼C, ∼DRB1, ∼DQA1, and ∼DQB1 Diversity in South African Populations. Frontiers in Genetics. 2022;13.

42. Jiang N, Yu Y, Zhang M, Tang Y, Wu D, Wang S, et al. Association between germ-line HLA and immune-related adverse events. Frontiers in Immunology. 2022;13.

43. Zaimoku Y, Patel BA, Adams SD, Shalhoub R, Groarke EM, Lee AAC, et al. HLA associations, somatic loss of HLA expression, and clinical outcomes in immune aplastic anemia. Blood. 2021;138(26):2799–809.

44. Arrieta-Bolaños E, Hernández-Zaragoza DI, Barquera R. An HLA map of the world: A comparison of HLA frequencies in 200 worldwide populations reveals diverse patterns for class I and class II. Frontiers in Genetics. 2023;14.

45. Lynch H. African Americans Chicago: Encyclopædia Britannica, Inc.; 2024 [updated Aug 19, 2024 cited 2024 23 August 2024]. Available from: https://www.britannica.com/topic/African-American.

46. Mungall AJ, Palmer SA, Sims SK, Edwards CA, Ashurst JL, Wilming L, et al. The DNA sequence and analysis of human chromosome 6. Nature. 2003;425(6960):805-11.

47. Sanjanwala B, Draghi M, Norman PJ, Guethlein LA, Parham P. Polymorphic Sites Away from the Bw4 Epitope That Affect Interaction of Bw4+ HLA-B with KIR3DL11. The Journal of Immunology. 2008;181(9):6293–300.

48. Augusto DG, Lobo-Alves SC, Melo MF, Pereira NF, Petzl-Erler ML. Activating KIR and HLA Bw4 Ligands Are Associated to Decreased Susceptibility to Pemphigus Foliaceus, an Autoimmune Blistering Skin Disease. PLOS ONE. 2012;7(7):e39991.

49. Tikly M, Njobvu P, McGill P. Spondyloarthritis in Sub-Saharan Africa. Current Rheumatology Reports. 2014;16(6):421.

50. Cauli A, Shaw J, Giles J, Hatano H, Rysnik O, Payeli S, et al. The arthritis-associated HLA-B*27:05 allele forms more cell surface B27 dimer and free heavy chain ligands for KIR3DL2 than HLA-B*27:09. Rheumatology (Oxford). 2013;52(11):1952–62.

51. Akkoç N, Yarkan H, Kenar G, Khan MA. Ankylosing Spondylitis: HLA-B*27-Positive Versus HLA-B*27-Negative Disease. Current Rheumatology Reports. 2017;19(5):26.

52. (SAA) SAoA. How Disease Severity, Ethnicity, and HLA-B27 Prevalence Intersect United State of America: Spondylitis Association of America (SAA); 2020 [Available from: https://spondylitis.org/spondylitis-plus/how-disease-severity-ethnicity-and-hla-b27-prevalence-intersect/.

53. Kopplin LJ, Mount G, Suhler EB. Review for Disease of the Year: Epidemiology of HLA-B27 Associated Ocular Disorders. Ocul Immunol Inflamm. 2016;24(4):470–5.

54. Ørsted M, Hoffmann AA, Sverrisdóttir E, Nielsen KL, Kristensen TN. Genomic variation predicts adaptive evolutionary responses better than population bottleneck history. PLOS Genetics. 2019;15(6):e1008205.

55. Robertson A, Charlesworth D, Ober C. Effect of inbreeding avoidance on Hardy-Weinberg expectations: examples of neutral and selected loci. Genet Epidemiol. 1999;17(3):165–73.

56. Mack SJ, Gourraud P-A, Single RM, Thomson G, Hollenbach JA. Analytical Methods for Immunogenetic Population Data. In: Christiansen FT, Tait BD, editors. Immunogenetics: Methods and Applications in Clinical Practice. Totowa, NJ: Humana Press; 2012. p. 215-44.

57. Siewert KM, Voight BF. Detecting Long-Term Balancing Selection Using Allele Frequency Correlation. Mol Biol Evol. 2017;34(11):2996–3005.

58. Delph LF, Kelly JK. On the importance of balancing selection in plants. New Phytologist. 2014;201(1):45–56.

59. Ellegren H, Galtier N. Determinants of genetic diversity. Nature Reviews Genetics. 2016;17(7):422–33.

60. Barreiro LB, Laval G, Quach H, Patin E, Quintana-Murci L. Natural selection has driven population differentiation in modern humans. Nature Genetics. 2008;40(3):340–5.

61. Nonić M, Šijačić-Nikolić M. Genetic Diversity: Sources, Threats, and Conservation. In: Leal Filho W, Azul AM, Brandli L, Lange Salvia A, Wall T, editors. Life on Land. Cham: Springer International Publishing; 2021. p. 421-35.

62. Lewontin RC. The interaction of selection and linkage. I. General considerations; heterotic models. Genetics. 1964;49(1):49–67.

63. Castelli EC, Mendes-Junior CT, Veiga-Castelli LC, Pereira NF, Petzl-Erler ML, Donadi EA. Evaluation of computational methods for the reconstruction of HLA haplotypes. Tissue Antigens. 2010;76(6):459–66.

64. Gourraud P-A, Pappas DJ, Baouz A, Balère M-L, Garnier F, Marry E. High-resolution HLA-A, HLA-B, and HLA-DRB1 haplotype frequencies from the French Bone Marrow Donor Registry. Human Immunology. 2015;76(5):381–4.

65. Arnatkevičiūtė A, Fulcher BD, Fornito A. Chapter 14 - Uncovering the genetics of the human connectome. In: Schirmer MD, Arichi T, Chung AW, editors. Connectome Analysis: Academic Press; 2023. p. 309-41.

66. Lucek K, Willi Y. Drivers of linkage disequilibrium across a species’ geographic range. PLOS Genetics. 2021;17(3):e1009477.

67. Alter I, Gragert L, Fingerson S, Maiers M, Louzoun Y. HLA class I haplotype diversity is consistent with selection for frequent existing haplotypes. PLOS Computational Biology. 2017;13(8):e1005693.

68. Charlesworth B, Morgan MT, Charlesworth D. The effect of deleterious mutations on neutral molecular variation. Genetics. 1993;134(4):1289–303.

69. World Medical Association Declaration of Helsinki: ethical principles for medical research involving human subjects. Jama. 2013;310(20):2191-4.

70. Sutter A, Amberg A, Boyer S, Brigo A, Contrera JF, Custer LL, et al. Use of in silico systems and expert knowledge for structure-based assessment of potentially mutagenic impurities. Regulatory Toxicology and Pharmacology. 2013;67(1):39–52.

71. Milius RP, Mack SJ, Hollenbach JA, Pollack J, Heuer ML, Gragert L, et al. Genotype List String: a grammar for describing HLA and KIR genotyping results in a text string. Tissue Antigens. 2013;82(2):106–12.

72. Lancaster AK, Single RM, Mack SJ, Sochat V, Mariani MP, Webster GD. PyPop: a mature open-source software pipeline for population genomics. Frontiers in Immunology. 2024;15.

73. Sinnwell J, Schaid D. Haplo Stats User Manual Version 1.1.0 Statistical Methods for Haplotypes when Linkage Phase is Ambiguous. 2010.

74. Excoffier L, Lischer H. ARLEQUIN suite ver 3.5: a new series of programs to perform population genetics analyses under Linux and Windows. Molecular ecology resources. 2010;10:564–7.

75. Coombs JA, Letcher BH, Nislow KH. create: a software to create input files from diploid genotypic data for 52 genetic software programs. Mol Ecol Resour. 2008;8(3):578–80.

76. Guo SW, Thompson EA. Performing the exact test of Hardy-Weinberg proportion for multiple alleles. Biometrics. 1992;48(2):361–72.

77. Hedrick PW. Gametic Disequilibrium Measures: Proceed With Caution. Genetics. 1987;117(2):331–41.

78. Cramér H. Mathematical Methods of Statistics. Princeton: Princeton University Press; 1946.

79. Begovich AB, McClure GR, Suraj VC, Helmuth RC, Fildes N, Bugawan TL, et al. Polymorphism, recombination, and linkage disequilibrium within the HLA class II region. J Immunol. 1992;148(1):249–58.

80. Slatkin M. A correction to the exact test based on the Ewens sampling distribution. Genet Res. 1996;68(3):259–60.

81. Slatkin M. An exact test for neutrality based on the Ewens sampling distribution. Genet Res. 1994;64(1):71–4.

82. Benjamini Y, Hochberg Y. Controlling the False Discovery Rate: A Practical and Powerful Approach to Multiple Testing. Journal of the Royal Statistical Society: Series B (Methodological). 2018;57(1):289–300.

83. Whittaker RH. Vegetation of the Siskiyou Mountains, Oregon and California. Ecological Monographs. 1960;30(3):279–338.

84. Whittaker RH. Evolution and measurement of species diversity. Taxon. 1972;21:213–51.

85. Shannon CE. A Mathematical Theory of Communication. Bell System Technical Journal. 1948;27(3):379–423.

86. Simpson EH. Measurement of Diversity. Nature. 1949;163(4148):688-.

87. Jost L. Entropy and diversity. Oikos. 2006;113(2):363–75.

88. Jaccard P. THE DISTRIBUTION OF THE FLORA IN THE ALPINE ZONE.1. New Phytologist. 1912;11(2):37-50.

